# qMaLioffG: A single green fluorescent protein FLIM indicator enabling quantitative imaging of endogenous ATP

**DOI:** 10.1101/2023.08.29.555275

**Authors:** Satoshi Arai, Hideki Itoh, Cong Quang Vu, Mizuho Nakayama, Masanobu Oshima, Atsuya Morita, Kazuko Okamoto, Satoru Okuda, Aki Teranishi, Madori Osawa, Yoshiteru Tamura, Shigeaki Nonoyama, Megumi Takuma, Toshinori Fujie, Satya Ranjan Sarker, Thankiah Sudhaharan, Taketoshi Kiya, E. Birgitte Lane, Tetsuya Kitaguchi

**Affiliations:** WPI Nano Life Science Institute, Kanazawa University, Kakuma-machi, Kanazawa, 920-1192, Japan; Skin Research Institute of Singapore, Agency for Science, Technology and Research, 8A Biomedical Grove, #06-06 Immunos, Singapore 138648; Division of Genetics, Cancer Research Institute, Kanazawa University, Kakuma-machi, Kanazawa 920-1192, Japan; Amphibian Research Center, Hiroshima University, Higashi-Hiroshima, 739-8526, Japan; Department of Pediatrics, National Defense Medical College, Saitama, Japan; School of Life Science and Technology, Tokyo Institute of Technology; Living Systems Materialogy (LiSM) Research Group, International Research Frontiers Initiative (IRFI), Tokyo Institute of Technology; Research Support Centre, Agency for Science, Technology and Research, 30 Biopolis Street, Matrix, Singapore 138671; Division of Life Sciences, Graduate School of Natural Science and Technology, Kanazawa University, Kakuma-machi, Kanazawa, 920-1192, Japan; Laboratory for Chemistry and Life Science, Institute of Innovative Research, Tokyo Institute of Technology, 226-8503, Kanagawa, Japan

**Keywords:** fluorescent protein probes, fluorescence lifetime imaging microscopy, ATP, live cell imaging, single cell analysis

## Abstract

The widespread use of fluorescence lifetime imaging microscopy (FLIM) for quantitative imaging is hindered by the limited availability of a FLIM-based genetically encoded indicator using a conventional 488 nm laser. Here, we present qMaLioffG, a single green fluorescent protein FLIM indicator showing a fluorescence lifetime change in ATP concentration within the physiological range. This allows quantitative imaging of endogenous ATP to investigate cellular energy status of different cell types.

---

Adenosine triphosphate (ATP) serves as the intracellular energy currency, driving energy metabolism and regulating extracellular signaling. Apart from these well-recognized functions, ATP has garnered attention for its role as hydrotrope, enhancing the solubility of intracellular proteins.^1^ To unravel biological processes involving ATP, fluorescence imaging has been indispensable for visualizing the dynamics of intracellular ATP. To date, various fluorescent ATP indicators have been developed, including Förster resonance energy transfer (FRET),^2^ dual-emission ratiometric,^3,4^ and single fluorescent protein (FP) based indicators.^5^ Recently, another newly developed indicator has emerged with submicromolar affinity for the detection of extracellular ATP.^6^ Despite the expanded availability, intensity-based analysis encounters common difficulties related to indicator concentration, excitation light amplitude, photobleaching and focus drift, which often hamper quantitative analysis.

Here, we present a single FP-based ATP indicator using fluorescence lifetime imaging microscopy (FLIM). Unlike intensity-based approach, FLIM based indicators offer the potential for quantitative analysis.^7^ For instance, FRET-FLIM utilizes FRET-based indicators where the fluorescence lifetime of the donor is altered during the energy transfer process. However, the use of a FRET indicator, which comprises two FPs, introduces uncertainties in quantitative analysis due to the variation in chromophore maturation states between the two proteins.^3^ In addition, FRET-FLIM microscopy occupies two color channels, which restricts the possibility for multiplex imaging. In contrast, a single FP-based FLIM indicator can potentially circumvent these issues. Notably, several indicators using a single FP like mTurquoise or T-Sapphire have recently emerged for detecting calcium^8^, glucose^9^, and lactate.^10^ Moreover, among several red-colored indicators, one specific red-colored FP-based Ca^2+^ indicator, R-CaMP1h, exhibits a capability for FLIM imaging.^11^ However, the availability of FLIM-based indicators compatible with a conventional 488 nm laser remains scarce, except the H_2_O_2_ indicator.^12^ In this study, we engineered a single green FP-based FLIM indicator with a high dynamic range for quantification of intracellular ATP. Using this indicator with a 488 nm pulsed laser, we conducted quantitative FLIM imaging of ATP in different cell types. Furthermore, we validated this indicator in multicellular systems including cultured HeLa cell spheroids and the *Drosophila* brain.

We previously reported MaLionG, an intensiometric ATP indicator based on single green FP.^5^ This was identified by generation of mutants where an ε subunit of an ATP-binding domain of a bacterial F_o_F_1_-ATP synthase was inserted into a variant of single green FP (Citrine) via peptide linkers (Fig. 1A). By varying the position of the binding domain inserted into Citrine and its peptide linkers, we optimized MaLionG with “turn-on” properties, i.e. exhibiting increased fluorescence intensity in the presence of ATP. Additionally, we also found another variant where the fluorescence intensity (λ_ex_/λ_em_ = 512/525) declined by 65% in the presence of 10 mM ATP (Fig. 1B). Intriguingly, while the fluorescence lifetime of MaLionG increased by only 0.16 ns, it decreased by 1.1 ns under 10 mM ATP, which was designated qMaLioffG (<u>q</u>uantitative <u>m</u>onitoring <u>A</u>TP <u>l</u>evel fluorescence l<u>i</u>fetime-based turn-<u>off</u> green) (Fig. 1C, Supplementary Fig. 1). The dynamic range (Δτ) of qMaLioffG was similar to that of mTurquoise2 based FLIM indicator (1.3 ns)^8^ but greater than that of conventional FRET-FLIM indicators (0.1–0.6 ns)^13^. The changes in fluorescence intensity and lifetime exhibited ATP-dose dependence and the *K*_d_ value was estimated to be 2.2 mM (Fig. 1E, Supplementary Fig. 2). In addition, the fluorescence intensity of qMaLioffG declined with decreasing pH, a typical characteristic observed in green FP indicators. However, the influence of pH fluctuation on fluorescence lifetime was observed to be less significant compared to its effect on the fluorescence intensity (Supplementary Fig. 3). Similar to MaLionG, qMaLioffG also showed specificity to ATP over other nucleotides (Supplementary Fig. 4). To investigate the sensing mechanism, we examined the absorption spectra in MaLionG and qMaLioffG (Fig. 1D). The absorbance of MaLionG displayed two peaks at 423 nm and 500 nm, corresponding respectively to protonated and deprotonated states of the tyrosine-based chromophore. In the presence of ATP, the 423 nm peak decreased while the 500 nm peak increased, resulting in the fluorescence enhancement. This was attributed to the deprotonated anionic species having a higher quantum yield than the protonated one, which is a common feature of a tyrosine-based single FP indicator. Similarly, the absorbance of qMaLioffG also exhibited two peaks at 417 and 505 nm. However, the absorbance change at 417 nm upon ATP binding was negligible while the other peak slightly shifted from 505 to 494 nm with small change in absorbance. It appears that qMaLioffG is unlikely to follow the exchange mechanism between protonated and deprotonated states (to be discussed later). Subsequently, we evaluated the performance of qMaLioffG as a FLIM indicator in HeLa cells. When qMaLioffG was expressed in HeLa cells, the application of sodium fluoride (NaF) as an inhibitor of enolase in glycolysis increased the fluorescence intensity and lifetime, indicating that it could detect the depletion of intracellular ATP levels (Fig. 1F– H, Supplementary Fig. 5–7). Intracellular ATP was quantified using calibration curves of fluorescence lifetime of pure qMaLioffG against ATP concentration using membrane-permeabilized cells as well as in solution (Fig. 1E).

**Figure 1.**
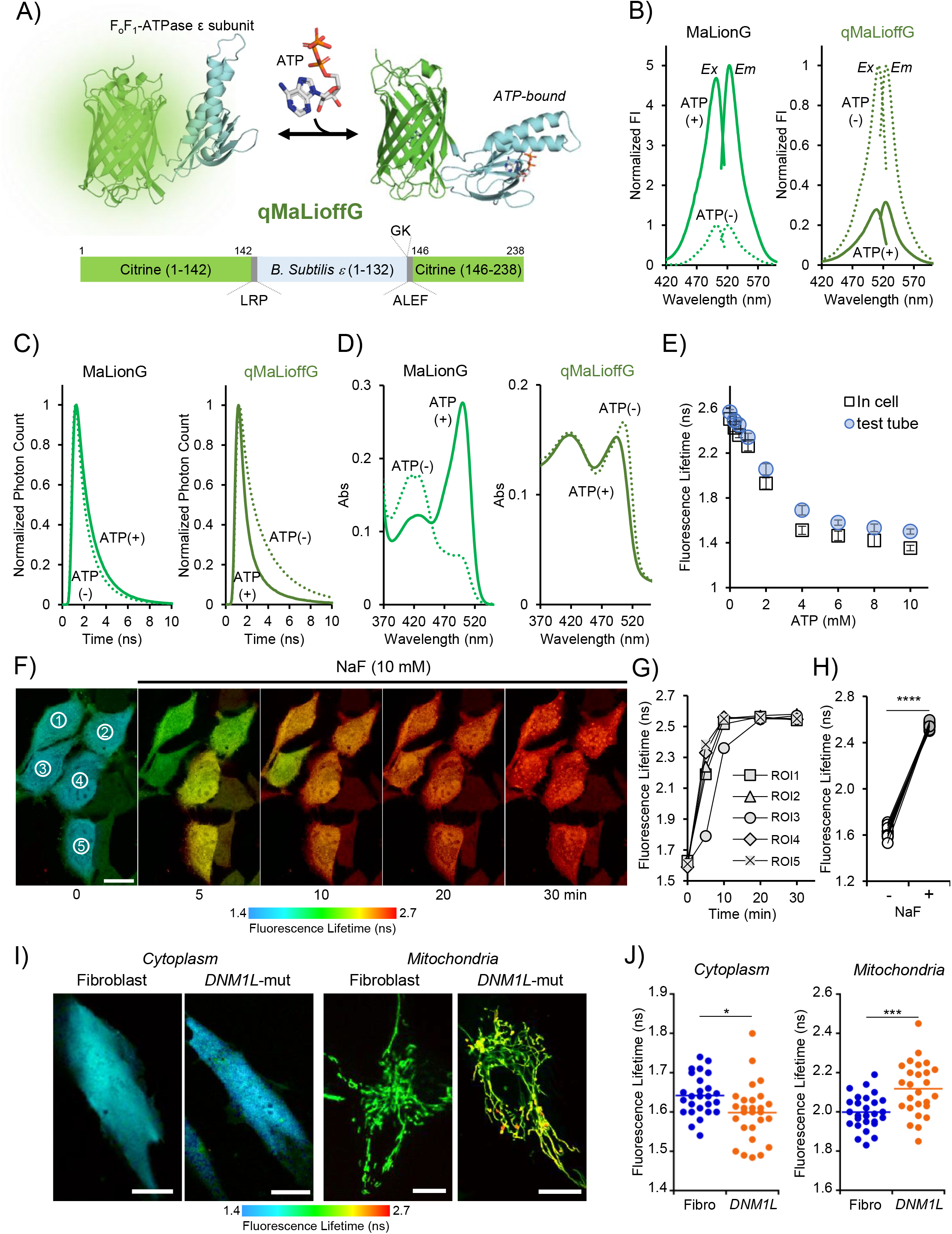
Characterization and validation of qMaLioffG. A) Schematic image of qMaLioffG, illustrated using AlphaFold2. B–D) Fluorescence spectra (B), fluorescence lifetime (C) and absorption spectra (D) of qMaLioffG and MaLionG under buffer conditions. The dotted lines represent the absence of ATP while solid lines represent the presence of 10 mM ATP. Fig. 1C: fluorescence lifetime values of MaLionG (τ = 1.39 ns and 1.56 ns) and qMaLioffG (τ = 2.57 ns and 1.49 ns) without and with ATP respectively. E) Calibration curves in the buffer solution and membrane-permeabilized HeLa cells. F–H) Validation of HeLa cells expressing qMaLioffG under NaF inhibition (10 mM). The montage (F), the time course of fluorescence lifetime at different cells (G), and the comparison before and 30 min after NaF treatment (n = 15, three independent dishes, paired student-t test) were shown in the inhibition experiments. ****p< 0.0001, Scale bar: 50 μm. I–J) Comparison of cytoplasmic and mitochondrial ATP level in human dermal fibroblasts and the cells derived from human patient with *DNM1L* mutation (n = 26–27, unpaired student t-test). *p<0.05, ***p<0.001, Scale bar: 10 μm.

We sought to examine whether qMaLioffG could differentiate ATP levels using different cells with mitochondrial dysfunction or distinct activities in energy metabolism. Additionally, for monitoring mitochondrial ATP, we constructed mitochondria-target qMaLioffG and validated its performance through an oligomycin inhibition experiment (Supplementary Fig. 8–9). Firstly, qMaLioffG was applied to skin fibroblasts derived from a patient with known mitochondrial dysfunction. We used fibroblasts obtained by skin biopsy from a patient with a mutation in dynamin-1-like protein (*DNM1L*). Correct fission and fusion processes are essential mitochondrial functions; mutation in *DNM1L* leads to disruption of mitochondrial fission and is likely to result in an insufficient supply of ATP.^14^ As predicted, mitochondrial ATP levels were significantly lower in mutant cells than in control human dermal fibroblasts (Fig. 1 I–J). Notably, the diseased cell exhibited higher cytoplasmic ATP levels than the control, indicating a potential compensatory mechanism for ATP production through glycolysis to counteract the ATP depletion. We next investigated cytoplasmic and mitochondrial ATP in mouse embryonic stem cells (mESC). mESCs can maintain a naïve state in culture by adding 2i (MEK inhibitor and GSK3 inhibitor) and leukemia inhibitory factor (LIF)^15^. After removal of 2i and LIF (hereafter 2iLIF), expression of Nanog, an essential factor for sustaining naïve state^16^, was entirely lost while there was a slight presence of Oct3/4 expression, serving as a pluripotency marker (Supplementary Fig. 10). Subsequent quantitative ATP analysis of mESC showed that cytoplasmic ATP levels were higher in the presence of LIF than in its absence, while no significant difference was observed in mitochondrial ATP levels (Fig. 2A). According to previous literature, pluripotency is maintained in the presence of 2iLIF through combination with glycolysis and oxidative phosphorylation (OXPHOS), referred to as naïve pluripotent stem cells^17^. Conversely, in the absence of 2iLIF, it has also been suggested that ATP production relies solely on glycolysis without the assistance of OXPHOS. This hypothesis was consistent with the observed decrease in cytoplasmic ATP levels upon 2iLIF removal (Fig. 2A). Additionally, it could also be assumed that mitochondrial ATP levels remain lower due to the rapid efflux of ATP from the mitochondria into the cytoplasm, thereby hindering the detection of a significant difference.^2^

**Figure 2.**
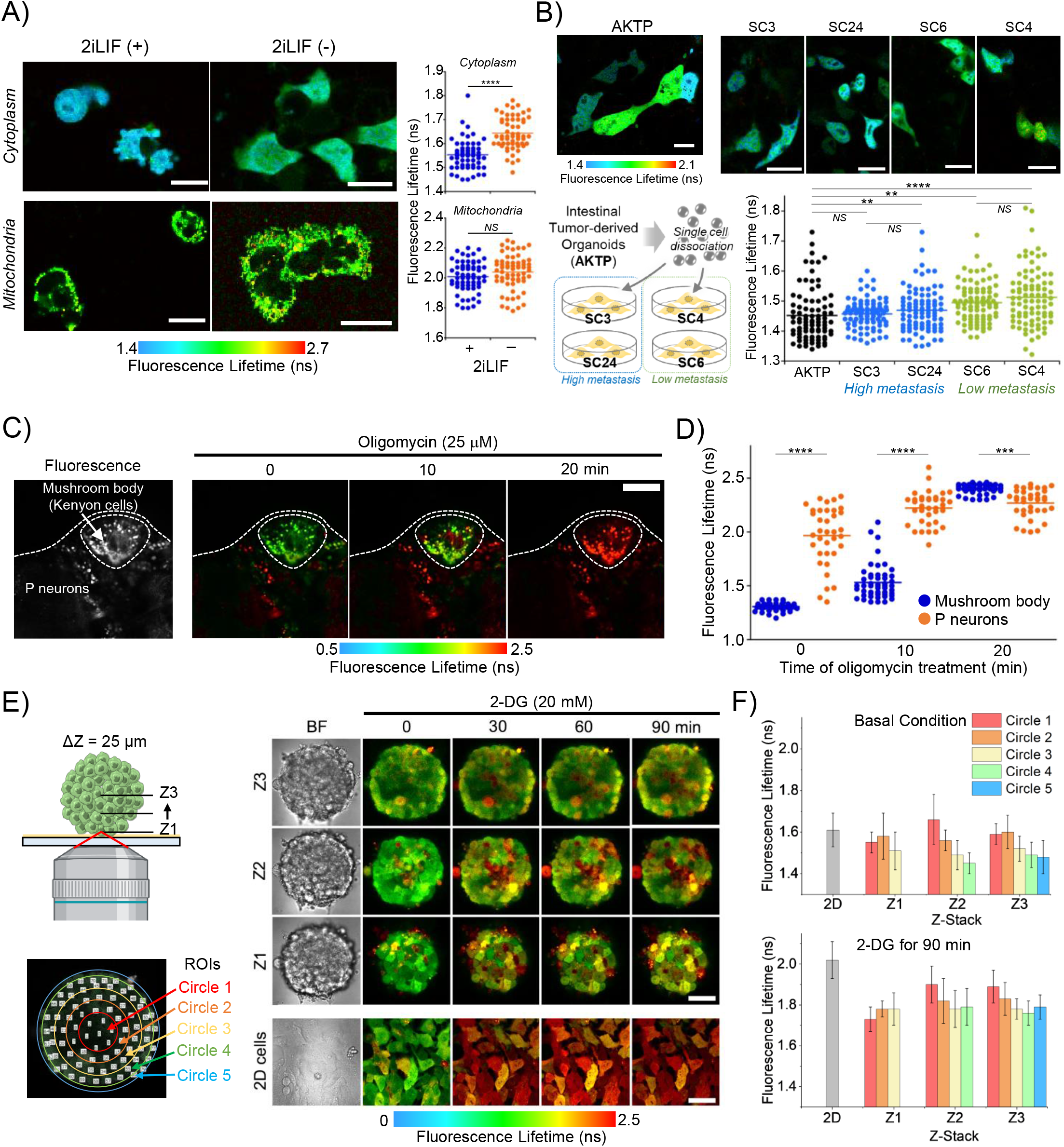
Applications for quantitative ATP mapping. A) Quantitative mapping of cytoplasmic and mitochondrial ATP in mouse embryonic stem cells (mESCs) with or without leukemia inhibitory factor (2iLIF) (n = 57–65, unpaired student t-test). Scale bar: 20 μm. ****p< 0.0001. NS (not significance). B) Analysis of cytoplasmic ATP in cells from intestinal tumor derived organoids (AKTP). Cell lines such as SC3, SC24, SC6 and SC4 were obtained via subcloning from AKTP, which were classified into two groups with high and low metastasis ability (n = 85–91, Tukey’s multiple comparisons). ****p<0.0001, **p<0.01, NS (not significance). The middle bar represents average. Scale bar: 20 μm. C–D) Visualization of the heterogeneity of ATP level in *Drosophila* brain. Fluorescence (left panel) and FLIM images of *Drosophila* brain under treatment with oligomycin (25 µM). (D) Quantification of ATP level between the mushroom body (Kenyon cells) and P neurons to the time of oligomycin treatment (n = 36–50, Tukey’s multiple comparisons). ****p<0.0001, ***p<0.001, Scale bar: 50 μm. E–F) Quantification of changes in ATP level under treatment of 2-DG in 3D spheroid and 2D HeLa cells stably expressing qMaLioffG. Schematic illustration of the imaging with definition of region of interests (ROIs) in 3D spheroid (E, left panel). FLIM images of 3D spheroid in different Z positions and 2D HeLa cells under the treatment of 2-DG (E, right panel). (F) Fluorescence lifetime of different ROIs in different Z positions at the basal condition and under the treatment of 2DG (20 mM) for 90 min. Data shows mean ± SD. Scale bar, 50 μm.

For more comprehensive understanding of proliferation and metastasis in cancer biology, quantitative imaging of cellular ATP offers valuable insights. Thus, we examined ATP levels in cancer cells with varying metastatic abilities. Previously, Morita et al. subcloned organoids derived from intestinal metastatic tumor carrying driver mutations such as *Apc*^*Δ716*^, *Kras*^*G12D*^, *Tgfbr2* ^*-/-*^ and *Trp53*^*R270H*^ (AKTP). They then transplanted a single subcloned cell line into mice to investigate the occurrence of liver metastasis.^18,19^ Subclones were thereafter categorized into two populations based on high (lines SC3 and SC24) or low (SC4 and SC6) metastatic abilities. Interestingly, we observed that cytoplasmic ATP levels were significantly higher in high metastasis cell lines than in low metastasis cell lines. No significant difference was observed for mitochondrial ATP (Fig. 2B, Supplementary Fig. 11). Further genetic analysis of these cell lines demonstrated the loss of stemness makers in cell lines with low metastatic ability, while the genetic background identified by AKTP remained unaltered.^19^ The observed higher ATP levels in metastatic cell lines corresponds well with the observation of elevated cytoplasmic ATP levels in mESC, which maintain stemness, as shown in Fig. 2A.

To further evaluate its performance of qMaLioffG in multicellular systems, it was applied to the *Drosophila* brain. Through bright field observation, the mushroom body and optic lobe were anatomically identified. FLIM imaging of the brain expressing qMaLioffG exhibited higher cytoplasmic ATP levels in the Kenyon cells of the mushroom body compared to the other neurons (P neurons, including pSP and pIP neurons) (Fig. 2C). This observation implies that the mushroom body, known for its pivotal role in learning and memory processes, likely necessitates a higher energy supply. Additionally, although ATP levels displayed variation among cells within the same region, the addition of oligomycin to the biospecimen significantly reduced the variance in ATP levels (Fig. 2C–D). In future studies, addressing such cellular heterogeneity could be accomplished by integrating other technologies that identify cell types based on genetic information.

Lastly, we attempted to image 3D spheroidal HeLa cells stably expressing qMaLioffG. In recent years, cultured spheroids have been used for drug screening as their denser multicellular structure is in some ways closer to tissues. Remarkably, we observed a lower cytoplasmic ATP level inside the spheroid compared to its surroundings (Fig. 2E–F). Although a systematic study has yet to be conducted, such a gradient in ATP concentration was rarely observed in smaller spheroid (data not shown), as well as 2D cell cultures. This observation is likely due to limited nutrient diffusion to the center of the spheroid. Subsequently, the spheroid was exposed to 2-deoxy glucose (2DG), a glycolytic inhibitor and a potential drug candidate. Over time, the concentration gradient of cytoplasmic ATP disappeared gradually, yet the heterogeneous ATP level persisted in comparison with 2D cells.

In this paper, we generated a first single green FP-based FLIM indicator and highlighted the potential applications using quantitative imaging of ATP. For reference, the fluorescence lifetime data obtained through this study was converted to ATP concentration using an in-cell calibration curve, whereupon it was seen to be in good agreement with the literature (Supplementary Table 1).^20^ Still, the exact sensing mechanism of the indicator underlying the alteration of fluorescence lifetime remains unclear. We hypothesize that ATP binding might increase the freedom of rotation of the free-moving bond, which connects the phenolic moiety and the imidazolidone in the tyrosine-based chromophore. As a result, while the deprotonated state of the chromophore is maintained to some extent, the energy from the excited state would be dissipated via a non-radiative process due to the acceleration of the chromophore motion. Supporting this hypothesis, we observed a significant change in the non-radiative rate constant (*k*_nr_) of qMaLioffG after ATP binding, while the change in the radiative rate constant (*k*_f_) was minimal (Supplementary Table 2). Consequently, the alteration of fluorescence lifetime is expected as it is inversely proportional to the combined kinetic rates of radiative and non-radiative processes.^7,21^ In the case of MaLionG, however, the persistent dominance of this anionic chromophore as the emitting species, with or without ATP, accounts for the observed negligible change in fluorescence lifetime. While a conventional FLIM system still suffers from relatively low temporal resolution, the photon-counting devices of the latest FLIM have been improved and allow imaging of one frame within a few hundred milliseconds to seconds.^22^ Single FP-based FLIM indicators will be a promising approach to provide us unique access to subcellular live cell physiology.

## Acknowledgements

We thank Dr. Taniyuki Furuyama (Kanazawa University) for kind support in measuring the fluorescence lifetime and quantum yield. This study was supported by the Japan Society for the Promotion of Science (JSPS) Grant-in-aid for Scientific Research (KAKENHI) (JP18H04832 and JP22H05176 to T.K.), a grant from the Nakatani Foundation (to S.A.), JST FOREST Program (JPMJFR201E to S.A. and JPMJFR203Q to T.F.), the Biomedical Research Council, Singapore through core support to the Skin Research Institute of Singapore (SRIS) (H.I., E.B.L.), a Human Frontier Science Program Grant (RGP0047/2018 to E.B.L.) (partial support for H.I.), and the Cooperative Research Program of “Network Joint Research Center for Materials and Devices” (to T.K. and S.A). We thank the A*STAR Microscopy Platform (AMP) for technical support. TS and the AMP are funded by Singapore’s Agency for Science, Technology & Research (A*STAR) through core funds and under the HBMS IAF-PP Project (H1701a0004) and through the National Research Foundation Singapore under its Shared Infrastructure Support grant for SingaScope – a Singapore-wide microscopy infrastructure network (NRF2017_SISFP10). We gratefully acknowledge financial support from the JSPS of Science Overseas Research Fellowships (to H.I.). We thank the Bloomington Stock Center for fly strains.

