## Supplementary information for "qMaLioffG: A single green fluorescent protein FLIM indicator enabling quantitative imaging of endogenous ATP"

Contents:

1) Supporting Figures

2) Experimental sections

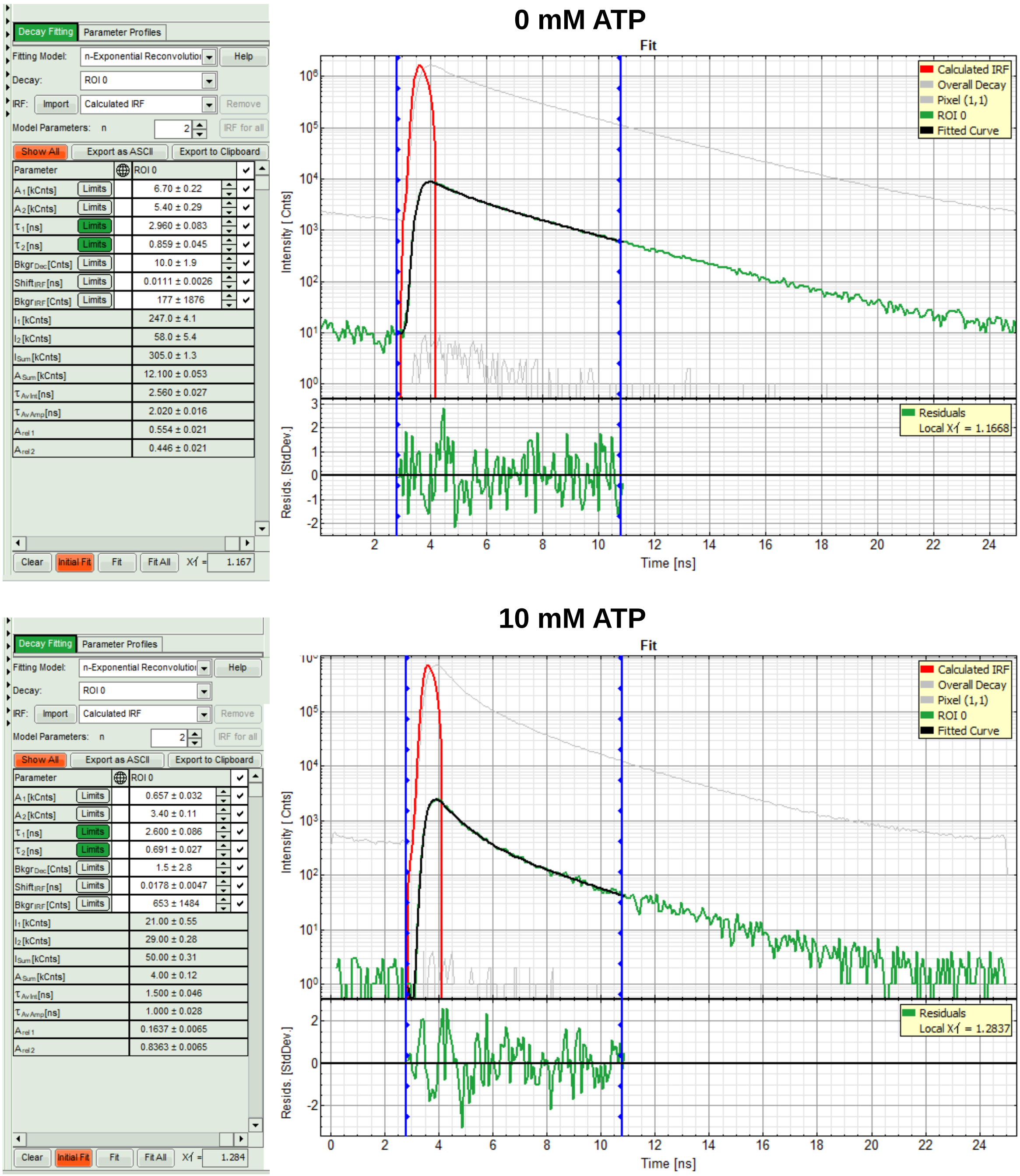

**Supplementary Figure 1.** Analysis of fluorescence lifetime. We performed fluorescence lifetime analysis using SymPhoTime 64 software from PicoQuant. We calculated the fluorescence lifetime using a standardized "tau8" value proposed by Yellen group (Díaz-García *et al., J. Neurosci. Res.* **97**, 946-960, doi:https://doi.org/10.1002/jnr.24433, 2019). Basically, we fit the fluorescence lifetime decay histogram with two-exponential function and the arrival time is fixed to 0-8 ns starting from the pulse laser (i.e., from 2.8 to 10.8 ns in this case). Finally, we reported the intensity-weighted average lifetime ($\tau_{\mathrm{Av\ Int}}$).

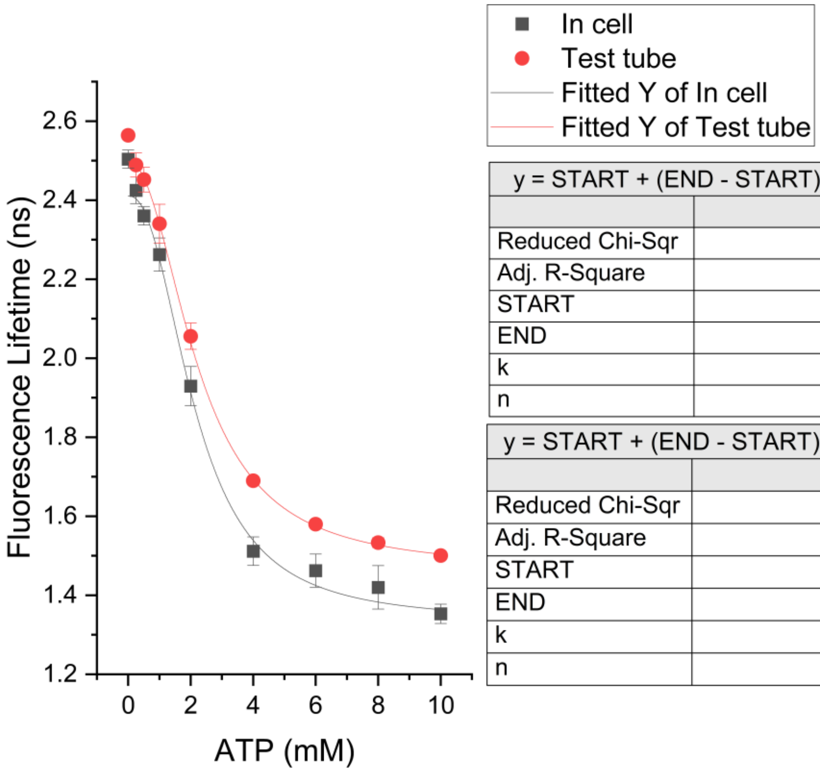

| y = START + (END - START) * x^n / (k^n + x^n) | |
|---|---|
| | In cell |
| Reduced Chi-Sqr | 3.90794 |
| Adj. R-Square | 0.98498 |
| START | 2.41076 ± 0.02818 |
| END | 1.33396 ± 0.03483 |
| k | 2.13817 ± 0.16031 |
| n | 2.30883 ± 0.36044 |

| y = START + (END - START) * x^n / (k^n + x^n) | |
|---|---|
| | Test tube |
| Reduced Chi-Sqr | 14.96245 |
| Adj. R-Square | 0.99125 |
| START | 2.49723 ± 0.02709 |
| END | 1.46408 ± 0.01478 |
| k | 2.22745 ± 0.10973 |
| n | 2.13676 ± 0.20282 |

**Supplementary Figure 2.** Curve fitting for ATP dose experiments. We performed curve fitting using modified Hill function with offset (Hill1 mode) in Origin software (OriginPro, Version 2022b. OriginLab Corporation, Northampton, MA, USA). The Hill1 equation is $y = START + (END - START)\frac{x^n}{k^n+x^n}$, where y is fluorescence lifetime, x is ATP concentration, k is dissociation constant, and n is the Hill coefficient.

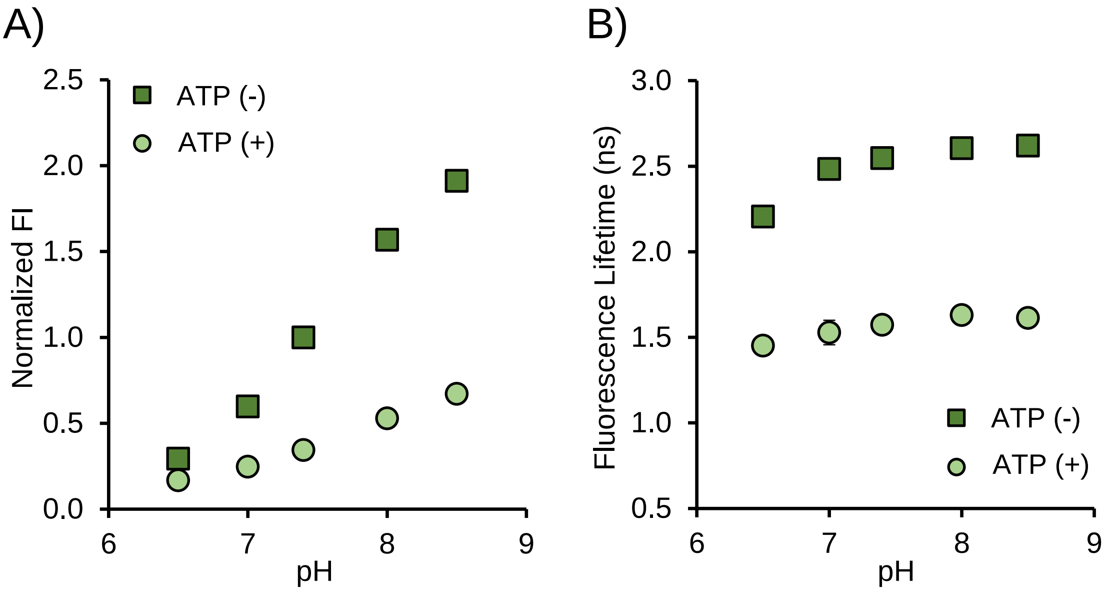

**Supplementary Figure 3**. pH sensitivity of qMaLioffG in the intensity-based (A) and FLIM-based analysis (B).

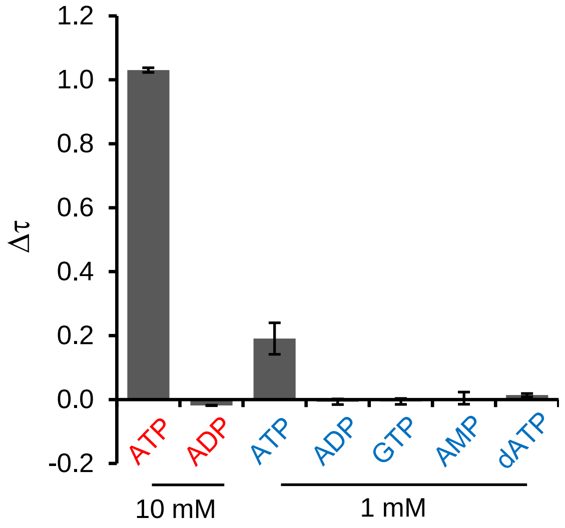

**Supplementary Figure 4**. Specificity of qMaLioffG to the other nucleotides. $\Delta\tau$ represents dynamic range obtained from the comparison with the presence and absence of nucleotides.

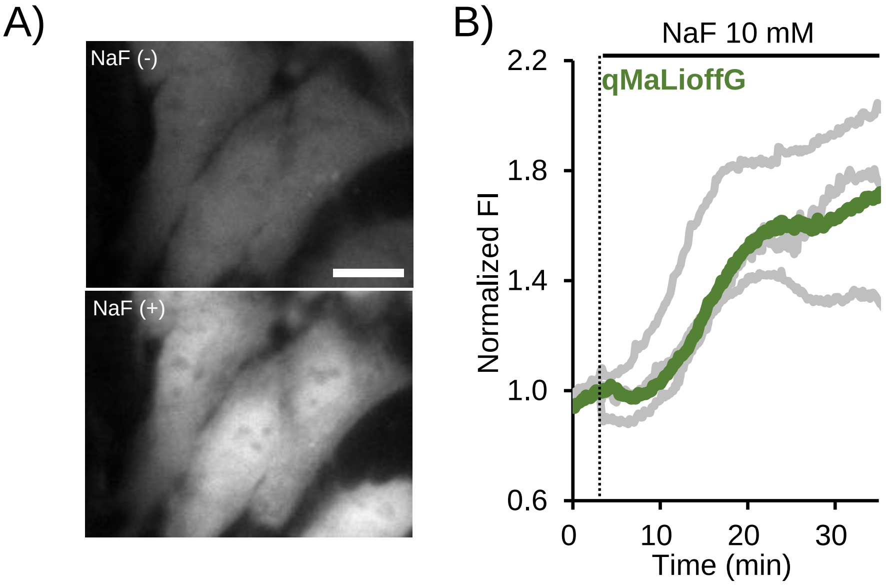

**Supplementary Figure 5**. Fluorescence intensity-based validation of qMaLioffG in HeLa cells in the inhibition experiments. A) Fluorescence image of HeLa cells expressing qMaLioffG before and after the treatment with NaF (10 mM). B) Average of normalized fluorescence intensity in single cells was evaluated per a dish (gray lines). Thick colour lines represent the average of three independent experiments (9 – 11 cells). Scale bar: 10 μm.

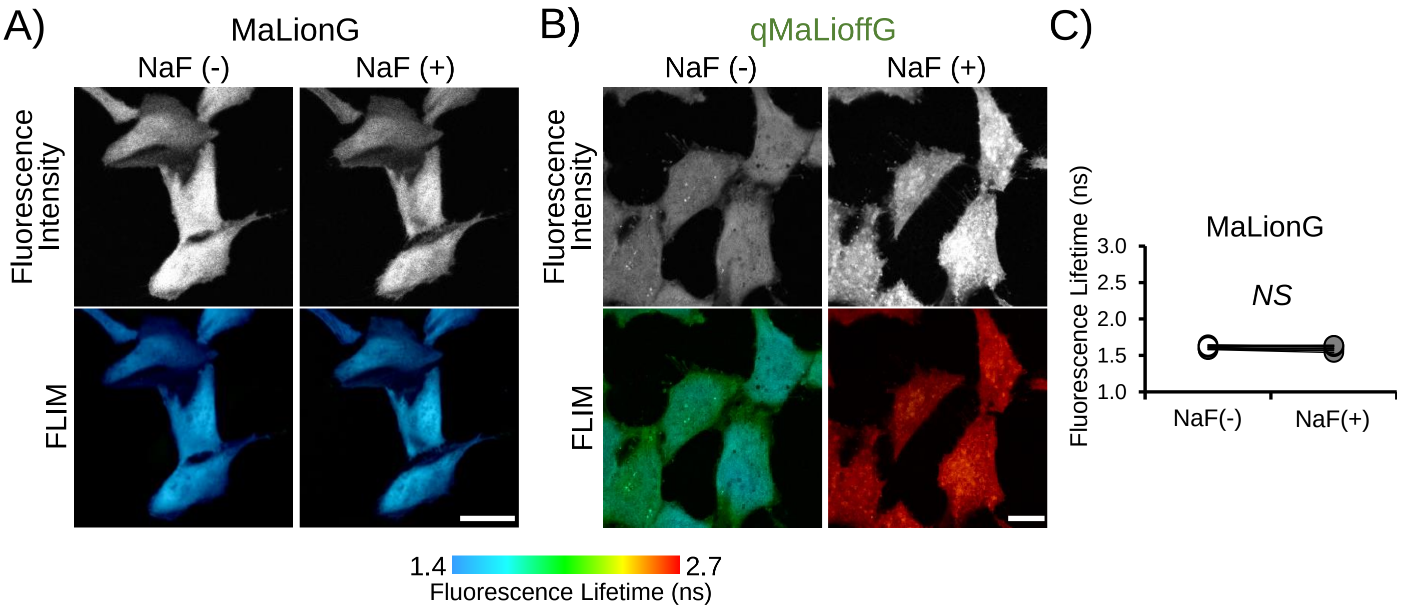

**Supplementary Figure 6**. Comparison with MaLionG and qMaLioffG in HeLa cells. A) Fluorescence intensity (upper panel) and FLIM (lower panel) images of MaLionG in the presence or absence of NaF (10 mM). B) Fluorescence intensity (upper panel) and FLIM (lower panel) images of qMaLioffG in the presence or absence of NaF (10 mM). Scale bar: 40 μm. C) The comparison before and 30 min after NaF treatment (n = 13, three independent dishes, paired student-t test). NS (not significance).

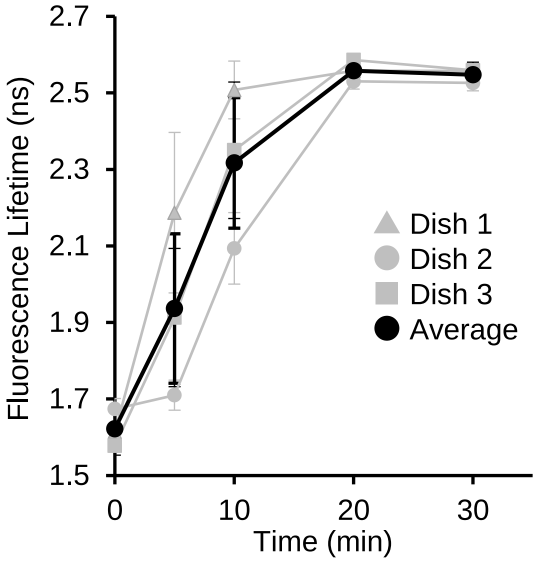

**Supplementary Figure 7**. Validation of HeLa expressing qMaLioffG under NaF inhibition experiments (10 mM). Same NaF inhibition experiment with Fig.1F-H were repeated in three independent dishes. The data showed means ± SD (n = 15).

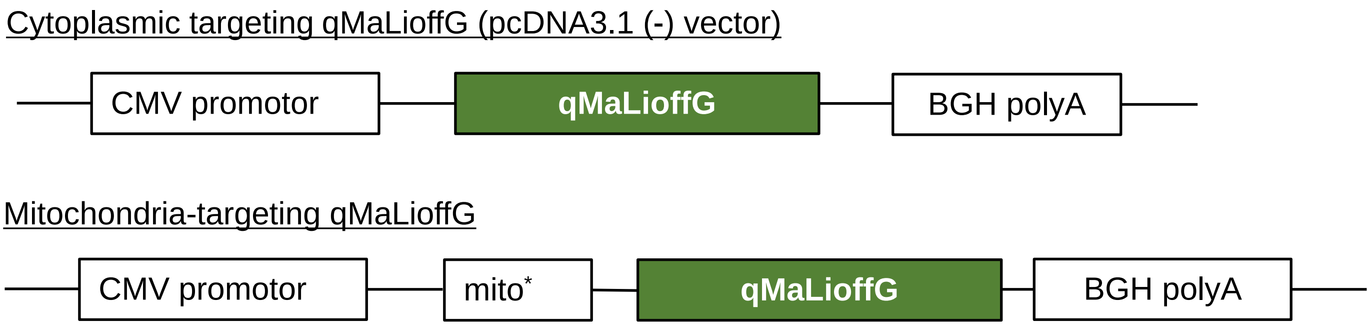

* Mitochondria targeting sequence; the subunit VIII of cytochrome c oxidase (SVLTPLLLRGLTGSARRLPVPRAKIHSL)

**Supplementary Figure 8**. Construction of plasmids for imaging experiments. Vectors for mammalians.

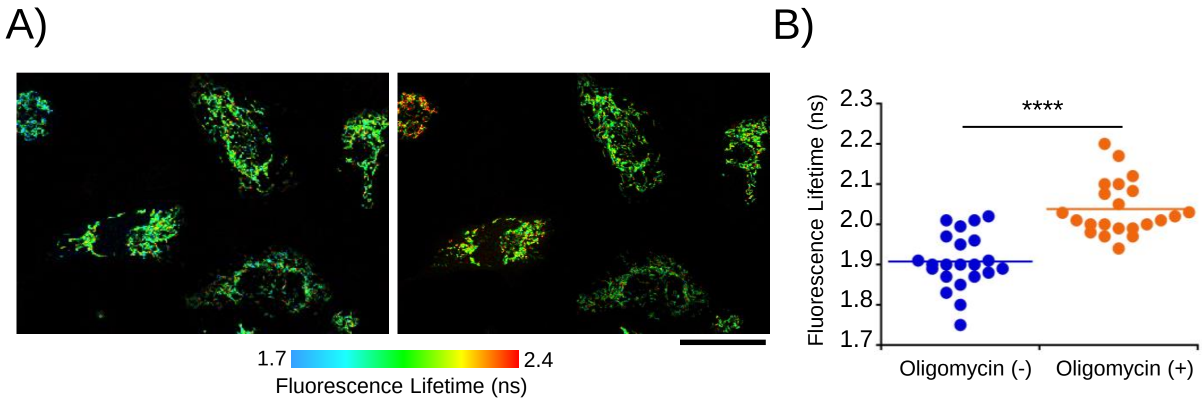

**Supplementary Figure 9**. Validation of mitoMaLioffG in HeLa cells. A) Mitochondrial ATP levels declined after the treatment with oligomycin (12.5 μM). B) Comparison of mitochondrial ATP level before and after the stimulation with oligomycin (n = 22, unpaired student t-test). Scale bar: 40 μm. ****$p<0.0001$

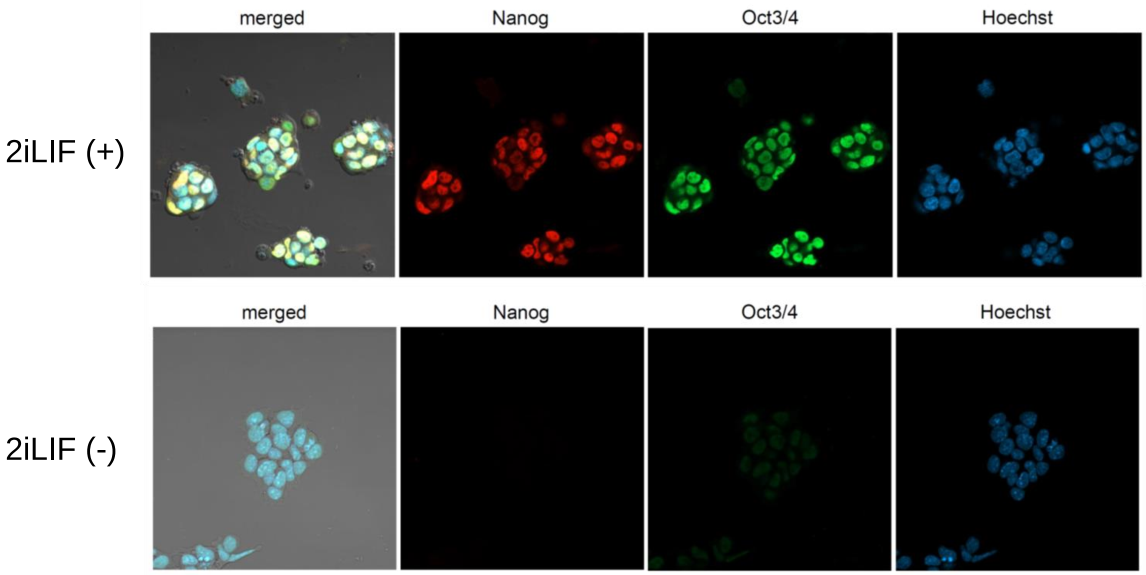

**Supplementary Figure 10.** Evaluation of expression level of Nanog and Oct3/4 as pluripotency markers in the presence or the absence of LIF. The both protein expression of Nanog and Oct3/4 in mouse ES cells in the presence or absence of LIF were evaluated using immunofluorescent staining. Nuclei were stained with Hoechst 33258.

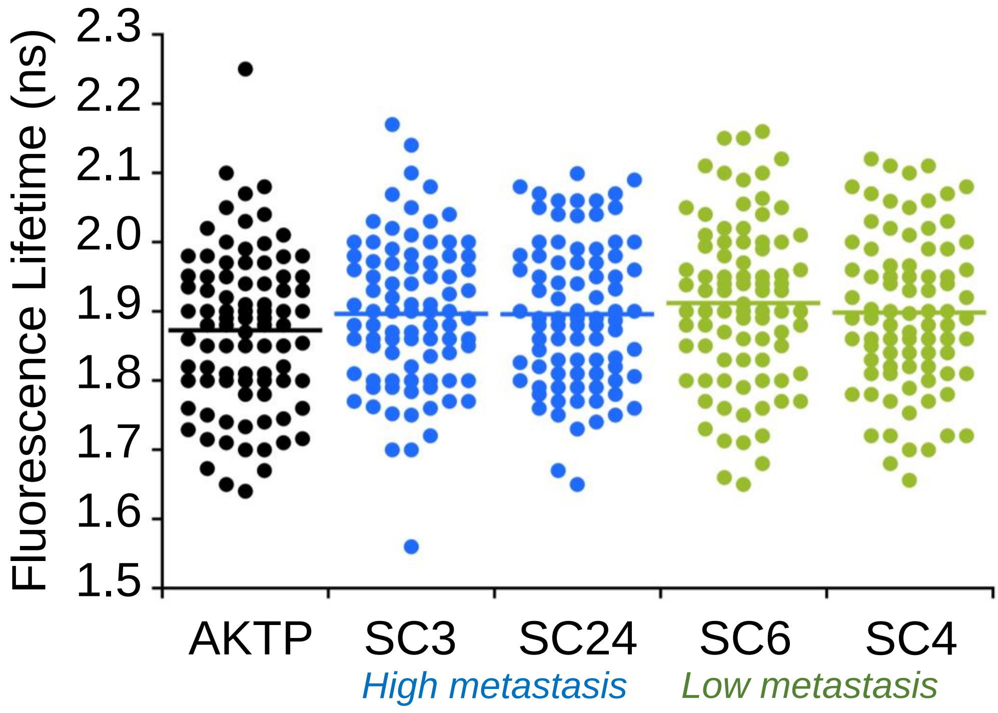

**Supplementary Figure 11**. Mitochondrial ATP in cells from intestinal tumor derived organoids (AKTP). Cell lines such as SC3, SC24, SC6 and SC4 were obtained via subcloning from AKTP, which were same with main Figure 2 (n = 85–88, Tukey's multiple comparisons). The significance in any combinations were not observed.

**Supplementary Table 1.** Evaluation of ATP level in all experiments

| | Cytoplasm [mM] | Mitochondria [mM] |
|---|---|---|
| ***Mamamalian cells*** | | |
| HeLa | 3.4 ± 0.3 | 2.1 ± 0.3 |
| Human epidermal fibloblast | 3.3 ± 0.3 | 1.8 ± 0.3 |
| mutated DNM1L | 3.6 ± 0.5 | 1.3 ± 0.5 |
| mES (with LIF) | 3.9 ± 0.5 | 1.7 ± 0.4 |
| mES (without LIF) | 3.3 ± 0.4 | 1.6 ± 0.4 |
| AKTP (tumor) | 5.3 ± 1.1 | 2.3 ± 0.5 |
| SC3 (with high metastasis) | 5.0 ± 0.8 | 2.2 ± 0.4 |
| SC24 (with high metastasis) | 4.9 ± 0.9 | 2.2 ± 0.4 |
| SC6 (with low metastasis) | 4.6 ± 0.8 | 2.1 ± 0.5 |
| SC4 (with low metastasis) | 4.5 ± 1.0 | 2.2 ± 0.5 |
| ***Drosophila*** | | |
| Mushroom body | >6.5 (out of range) | |
| Mushroom body (oligomycin 10 min.) | 4.5 ± 1.3 | |
| Mushroom body (oligomycin 20 min.) | 0.4 ± 0.1 | |
| Optic lobe | 2.1 ± 1.4 | |
| Optic lobe (oligomycin 10 min.) | 1.0 ± 0.5 | |
| Optic lobe (oligomycin 20 min.) | 0.8 ± 0.4 | |

**Supplementary Table 2.** Photochemical parameters of qMaLioffG and MaLionG

| | | 0 mM ATP | 10 mM ATP |
|---|---|---|---|
| **qMaLioffG** | **QY** | 0.247 | 0.131 |
| | **Lifetime (ns)** | 2.51 | 1.24 |
| | **$k_r$ ($x10^8$; $s^{-1}$)** | 0.98 | 1.06 |
| | **$k_{nr}$ ($x10^8$; $s^{-1}$)** | 3.00 | 7.00 |
| **MaLionG** | **QY** | 0.142 | 0.318 |
| | **Lifetime (ns)** | 1.28 | 1.54 |
| | **$k_r$ ($x10^8$; $s^{-1}$)** | 1.11 | 2.06 |
| | **$k_{nr}$ ($x10^8$; $s^{-1}$)** | 6.72 | 4.42 |

**Chemicals**

For the test to check the specificity, ATP, ADP, AMP and GTP were purchased from Nacalai Tesque, and dATP was purchased from Thermo Scientific. For cellular studies, sodium fluoride (NaF) and oligomycin were purchased from Sigma-Aldrich. All oligonucleotides for primers were purchased from Sigma-Aldrich. Common reagents for cell culture were as follows: DMEM (Nacalai tesque), GlutaMax (Gibco), fatal bovine serum (Gibco), sodium pyruvate (Nacalai Tesque), penicillin-streptomycin (FujiFilm Wako), β-escin (FujiFilm, MP Biomedicals, Inc.), and EGTA (Sigma-Aldrich).

**Screening and characterization of qMaLioffG**

As mentioned in main manuscript, the screening procedure of qMaLioffG including protein purification and *in vitro* spectroscopic characterization were addressed in supporting information in previous paper (ref. 5 in main manuscript: Arai et al., Angew. Chemie Int. Ed, 2018). According to the previous paper, buffer solutions containing qMaLioffG (10 μM) and different nucleotides (ADP, AMP, GTP, dATP) were prepared and placed onto glass bottom dishes and fluorescence lifetime was evaluated using the microscopy. Likewise, the pH sensitivity of qMaLioffG was evaluated with varying pH (from 6.5 to 8.5). The in-cell calibration curve between ATP and fluorescence lifetime was prepared according to the previous paper.[1] Briefly, the buffer solution with K-Gluconate (140 mM), NaCl (10 mM), HEPES (10 mM), EGTA (1 mM), $MgCl_2$ (1.3 mM), and $CaCl_2$ (0.34 mM) was prepared, followed by the addition of β-escin as a cell-permeabilized reagent (adjusted to 50 μM as final concentration). After HeLa cells were treated with the solution for 5 min., it was exposed with the solution without β-escin. With the buffer solutions varying ATP concentration (0-8 mM), fluorescence lifetime was plotted.

**Fluorescence lifetime imaging microscopy (FLIM)**

FLIM imaging was performed using an FV1200 confocal microscope (Olympus) equipped with rapidFLIM$^{HiRes}$ with MultiHarp 150 Time-Correlated Single Photon Counting (TCSPC) unit (PicoQuant). An oil immersion objective lens was used through all experiments (PLAPON 60X, NA = 1.42, Olympus). For excitation, a 485 nm pulse laser (PicoQuant) was used, and fluorescence emission was collected through a bandpass filter (520/35 nm, Bright Line HC). The scanning size was 512 × 512 pixels, and the scanning time was 1.109 sec/frame. The analytical method was detailed in Figure S1 of supporting information.

**Fluorescence quantum yield and lifetime measurement**

The fluorescence quantum yield (QY) of purified qMaLioffG protein was measured with the Quantaurus-QY Plus UV-NIR absolute PL quantum yield spectrometer (C13534, Hamamatsu). The excitation wavelength was 480 nm. For fluorescence lifetime measurement, the Quantaurus-Tau Fluorescence lifetime spectrometer (C11367, Hamamatsu) was used. The excitation and emission were set to 470 and 525 nm, respectively. The peak counts were set to 500 and two-component fitting was performed to obtain the averaged $\tau$. The concentration of qMaLioffG was 10 μM, and the measurements were conducted in an ATP-binding buffer (50 mM MOPS, 50 mM KCl, 0.5 mM $MgCl_2$, 0.05 % Triton X-100, pH 7.4) for both fluorescence QY and lifetime measurements.

**Cell culture and imaging experiments for HeLa cells, human dermal fibroblasts, cells from the patient with *DNM1L*, and intestinal tumor-derived cell subclones.**

HeLa cells purchased from ATCC (CCL-2) were cultured in Dulbecco's modified Eagle's medium (DMEM) supplemented with fetal bovine serum (FBS, 10 %) and penicillin-streptomycin (1 %). Human dermal fibroblasts as a control were purchased from CELLnTEC (Bern, Switzerland). Fibroblasts were also established by skin biopsy from the patient with *DNM1L* mutation (a de novo missense mutation of c. 1217T>C, p.Leu406Ser).[2] These cells were cultured in DMEM with 10 % FBS and 1 % penicillin-streptomycin. The metastatic intestinal tumor-derived cell subclones were established previously.[3,4] In brief, metastatic intestinal tumor-derived organoids carrying $Apc^{\Delta716}$, $Kras^{G12D}$, $Tgfbr2^{-/-}$, $Trp53^{R270H}$ mutations (AKTP) were developed from intestinal tumors of AKTP mice.[3] Established AKTP organoid was subjected to subcloning, and metastatic ability of each subclone was examined by spleen transplantation and in vivo luciferase imaging. Although parental cells maintained high metastatic ability, approximately 30 % of subclones showed a loss of metastatic ability. In this study, SC3 and SC24 subclones were used as high metastatic AKTP tumor cells, while SC6 and SC4 were used as low or non-metastatic cells.

For further FLIM experiments, these cells were seeded on a 3.5 cm glass-based dish, followed by the transfection with 0.2 μg of the plasmid DNA of qMaLioffG (pcDNA3.1(-)) using 0.8 μl of FuGENE HD Transfection Reagent (Promega) in 10 μl of Opti-MEM (Life Technologies Corporation). After the transfection, the dishes were maintained at 37 °C under 5% $CO_2$ for 8 h, washed out with a fresh DMEM with 10 % FBS, and then incubated at 30 °C for 48 h before imaging experiments.

**ES cell culture and immunostaining**

A mouse ESC line EB5 (RIKEN Bioresource Center) were cultured under maintenance medium consisting high glucose DMEM, GlutaMax, 15 % fatal bovine serum, 1 mM sodium pyruvate, penicillin-streptomycin, non-essential amino acid (NEAA; Gibco), 0.1 mM 2-mercapthoethanol (Sigma-Aldrich), and 1000U/mL leukemia inhibitory factor (LIF; Merck/Sigma-Aldrich, ESG1107). Medium was filtrated with 0.22 µm filter unit (Corning). Cells were passaged on 0.1 % gelatin (Sigma-Aldrich)-coated plastic dishes (FALCON #353004) every other day. Before transfection of qMaLioffG (pcDNA3.1(-)), 1x $10^5$ cells were passaged on 3.5 cm glass bottom dishes (D11130H, Matsunami-Glass, Japan) with thin Matrigel (Corning, 354230) coating. Cells were incubated with maintenance medium with 2i (1 μM PD0325901 (Stemgent, USA, Stemolecule™ 04–0006) and 3 μM CHIR99021 (Stemgent, Stemolecule™ 04–0004)). 24 h after seeding, qMalioffG was transfected into cells by using FuGene

HD transfection reagent (Promega). Cells were incubated for 24 h, and then used for FLIM observation and fixed for immunostaining. For induction of differentiation, 2i and LIF were simultaneously removed from maintenance medium with transfection.

Cells were fixed with 4 % paraformaldehyde for overnight at 4 ºC. After fixation, cells were permeabilized with 0.5 % Triton-X diluted with PBS for 10 min at room temperature (RT). Cells were blocked with CAS-Block™ Histochemical Reagent (Thermofisher Scientific, 008120) for 30 min at RT. Then cells were incubated with primary antibodies against Nanog (abcam, ab80892) and Oct3/4 (Santa Cruz Biotechnology, sc-5279) for 1 h at RT. Cells were washed with PBS three times, and incubated with secondary antibodies, ab150116 and ab150080 (abcam). Nuclei were counterstained with 50 mg/ml Hoechst 33258 (Dojinkagaku, Japan). After incubation, cells were washed with PBS, and mounted with Fluoro-KEEPER antifade reagent (Nacalai tesque). Fluorescent images were collected with Zeiss LSM800 with x 40 lens (Plan-APO CHROMAT 40x/1.4 Oil DIC).

**Fly experiments**

Flies (*Drosophila melanogaster*) were maintained on standard fly medium, under normal 12-h light/12-h dark conditions at 25 °C. MaLioffG-fly, a codon-optimized MaLioffG gene for *Drosophila melanogaster* (Eurofingenomics, Japan), was inserted into the pJFRC7 (Addgene: Plasmid #26220) vector (Pfeiffer et al., 2010) via the *Xho* I and *Xba* I restriction sites. The plasmid was injected into embryos, which have nos-phiC31 (BL# 34771) integrase and VK00005 (BL# 9725), and the UAS-MaLioffG strain was established with standard methods. Through the GAL4/UAS system, MaLioffG was expressed in *fruitless*-expressing neurons (BL#66696) (Stockinger et al., 2005). Eclosed male flies were collected and maintained in a group (approximately 10-20 flies per vial) until use. In all experiments, flies 6-7 days after eclosion were used. The plasmid was obtained from Addgene (#26220).

All imaging experiments were performed in an adult hemolymph like (AHL) saline, containing 108 mM NaCl, 5 mM KCl, 2 mM $CaCl_2$, 8.2 mM $MgCl_2$, 4 mM $NaHCO_3$, 1 mM $NaH_2PO_4$, 5 mM trehalose, 10 mM sucrose, 5 mM HEPES (Wang et al., 2003). The brain was dissected in $Ca^{2+}$-free AHL saline, and the blood brain barrier was digested with papain (10 U/ml) (Worthington Biochemical Corporation, New Jersey, USA) for 15 min at room temperature. The brain was transferred into a glass bottom dish filled with AHL.

**Lenti viral infection**

For stable expression, lentivirus was used. 5 µg of CSII-EF-qMaLioffG, 4 µg of pMDLg/pRRE, 2 µg of pRSV-Rev, and 2.5 µg pMD2.G were transfected to HEK293T (seeded with 7 million cells/10 cm dish on a previous day) with 41 µL of Lipofectamine 3000. For the next two days after the transfection, the viral supernatant was collected with filtration each day and concentrated with Lenti-X Concentrator. HeLa cells were infected by the lentivirus with 10 µg/mL polybrene. Then, cells with positive expression were collected by BD FACSAria™ II Cell Sorter (Laser 488 nm, filter 530/30 for qMaLioffG).

**2-Deoxy-D-glucose (2DG) experiments of 3D spheroid and 2D HeLa cells**

We prepared 3D spheroid HeLa cells stably expressing qMaLioffG in a EZSPHERE 96-well plate (4860-900, Iwaki). We cultured HeLa cells with a density of $8x10^4$ cells/well in a Dulbecco's modified eagle's medium (DMEM, 11965092, ThermoFisher Scientific) supplemented with 10% fetal bovine serum (FBS) and 1% Penicillin-Streptomycin (P/S) at 37 °C in a 5% $CO_2$ incubator for 24–48 h to form spheroids. Next, we gently transferred spheroid cells from 96-well plate to a glass-bottom dishes coated with a functional polymer sheet[5] (Suematsu et al, J. Mater. Chem B, 8, 6999–7008 (2020)) and filled with 2 mL DMEM culture medium and continued cultured for 8-10 h. To remove unattached spheroid cells and dead cells, we washed the spheroid cells three times with PBS solution and finally exchanged the medium to Hanks' Balanced Salt solution (HBSS (+), 09735-75, Nacalai Tesque Inc.) before the microscopy observation. For 2DG treatment, we freshly prepared 2DG (D0051, TCI) and stimulated 3D spheroid or 2D HeLa cells stably expressing qMaLioffG with 2DG to the final concentration of 20 mM. We performed FLIM imaging with the same setup as mentioned in the FLIM imaging section otherwise stated.
